# Variable phage susceptibility of *Pseudomonas aeruginosa* from patients with and without cystic fibrosis following treatment-emergent resistance to ceftolozane-tazobactam

**DOI:** 10.64898/2026.09.03.749285

**Authors:** Nathan R. Wallace, Renata L. DiDonato, Blake A. Jackson, Ellen G. Kline, Kevin M. Squires, Ava J. Dorazio, Ryan K. Shields, Daria Van Tyne

**Affiliations:** Division of Infectious Diseases, University of Pittsburgh School of Medicine, Pittsburgh, Pennsylvania, USA; Center for Innovative Antimicrobial Therapy, University of Pittsburgh School of Medicine, Pittsburgh, Pennsylvania, USA; Center for Evolutionary Biology and Medicine, University of Pittsburgh School of Medicine, Pittsburgh, Pennsylvania, USA

**Keywords:** Antibiotic resistance, *P. aeruginosa*, phage therapy, cystic fibrosis

## Abstract

**Background:** *Pseudomonas aeruginosa* is a ubiquitous opportunistic bacterial pathogen associated with nosocomial infections and is a leading cause of infection in persons with cystic fibrosis (pwCF). The front-line treatment for multidrug-resistant *P. aeruginosa* infections is ceftolozane-tazobactam (C/T). While previous research has characterized clinical *P. aeruginosa* isolates that evolved resistance to C/T, the collateral effect of evolved resistance on susceptibility to bacteriophages has not been explored.

**Methods:** We collected paired *P. aeruginosa* clinical isolates from 10 pwCF and 18 non-pwCF who developed treatment-emergent C/T resistance. We compared genetic relatedness, acute and chronic virulence phenotypes, and antibiotic and phage susceptibilities between each pair of susceptible baseline and treatment-emergent C/T-resistant isolates.

**Results:** Treatment-emergent C/T-resistant isolates were genetically closely related to baseline isolates in all patients. Virulence phenotypes did not differ between pre-and post-C/T exposure isolates, but isolates from pwCF demonstrated differences in protease production, twitching motility, and amino acid auxotrophy compared to isolates from non-pwCF. Treatment-emergent C/T resistance was associated with increased resistance to ceftazidime and ceftazidime/avibactam, but no other trends in antibiotic or phage susceptibility were detected.

**Conclusions:** Treatment-emergent resistance to C/T does not cause predictable alterations in phage susceptibility across genotypically and phenotypically diverse multidrug-resistant *P. aeruginosa* clinical isolates.

## 1. Introduction

Antimicrobial development is far outmatched by the emergence of multidrug-resistant (MDR) bacteria (1–3). MDR *Pseudomonas aeruginosa* is a pervasive Gram-negative opportunistic pathogen that causes tens of thousands of infections in hospitalized patients and thousands of deaths (4–6). Persons with cystic fibrosis (pwCF) represent unique hosts for *P. aeruginosa*, whereby initial infection results in chronic respiratory tract colonization due to impaired mucociliary clearance followed by pathoadaptation over years to decades (7–11). Once chronic infections are established, antibiotic therapy is rarely able to eradicate them (7–11). Chronic bacterial infections in pwCF represent a significant cause of morbidity and mortality in this population (7–11).

The antibiotic arsenal available to treat *P. aeruginosa* infections is limited by the occurrence of MDR strains, some of which are resistant to all available antibiotics (12). To combat MDR *P. aeruginosa* infections, the beta-lactam/beta-lactamase inhibitor combination ceftolozane-tazobactam (C/T) was deployed starting in 2014 (5,12). Ceftolozane is a fifth generation oxyimino-cephalosporin with a chemical structure similar to ceftazidime, but harbors more favorable *P. aeruginosa* activity (12,13). Despite demonstrating favorable clinical outcomes, clinical and *in vitro* resistance to C/T has been observed (5,13,14). Prior studies have shown that mutations in *ampC*, *ampR*, *pilB*, *PA3206* (a putative two-component sensor), *clpX*, *mex* operons, and *acrB* are associated with C/T resistance (5,13,15,16).

A promising alternate strategy for treating MDR *P. aeruginosa* infections is bacteriophage (phage) therapy, which utilizes viruses that selectively infect and kill bacteria to treat infections (2,5–7,17). Phages are pervasive in the environment, and phage therapy takes advantage of the predator-prey relationship between phages and bacteria to target bacteria causing infections (18). The use of phages in conjunction with antibiotics could provide a two-pronged approach to treat bacterial infections, particularly those caused by MDR bacteria and in pwCF (19,20). Here we collected pairs of clinical *P. aeruginosa* isolates from 28 patients (including 10 pwCF) that were isolated before and after treatment-emergent resistance to C/T. We sequenced the genomes of all 56 isolates, assessed virulence-associated phenotypes, and evaluated susceptibility to 10 clinically available antibiotics and seven lytic *P. aeruginosa*-targeting phages. Combining the genomic and phenotypic data collected, we characterized patterns of antibiotic and phage susceptibility that could inform the treatment of pwCF with these therapies.

## 2. Methods

### 2.1 Bacterial isolate collection

Clinical bacterial isolates from bronchoalveolar lavage (BAL) fluid and sputum were collected from patients with MDR *P. aeruginosa* infections before and after C/T treatment. “Treatment-emergent C/T resistance” was defined as a ≥4-fold increase in the C/T minimum inhibitory concentration (MIC) between pre-exposure and post-exposure isolates (21). Isolates were speciated via MALDI-TOF at the UPMC Clinical Microbiology Lab, sub-cultured in Tryptic Soy Broth (TSB), and cryopreserved at −80°C in TSB with 16.7% glycerol. Bacterial cultures for experiments were started by inoculating 2 mL of TSB, Brain Heart Infusion (BHI) broth, or Lysogeny Broth (LB) from cryopreserved stocks. Isolate collection was approved by the University of Pittsburgh Institutional Review Board under STUDY22070065.

### 2.2 Whole genome sequencing and analysis

*P. aeruginosa* genomic DNA was extracted from 1 mL of overnight bacterial cultures grown in BHI using the DNeasy Blood & Tissue Kit (Qiagen, Germantown, MD) according to the manufacturer’s instructions. Short-read sequencing libraries were generated with a Nextera library preparation kit (Illumina, San Diego, CA). Libraries were sequenced on an Illumina MiSeq with 300-bp paired-end reads or on a NextSeq with 150-bp paired-end reads. For long-read sequencing, libraries were prepared using the ONT Rapid Barcoding Kit (SQK-RBK004; Oxford, UK) and sequenced on a MinION instrument. Nanopore sequencing data were basecalled using Guppy v4.4.0. Long-and short-read data were hybrid assembled with Unicycler (22). Genomes were annotated with prokka and a phylogenetic tree was constructed from a core genome alignment generated by Roary using RAxML with the GRTCAT algorithm and 100 bootstraps (23–25).

### 2.3 Virulence phenotyping

Protease production was assessed by spotting 3 uL overnight cultures grown in BHI broth onto BHI 1.5% agar plates containing 10% nonfat milk. Plates were incubated for 24 hours at 37°C and protease production was scored categorically by visual inspection (0 = no clearance; 1 = halo around colony; 2 = thin ring, full clearance; 3 = ring, full clearance; 4 = large ring, full clearance). Twitching motility was assessed by stabbing 10 uL pipette tips dipped into overnight cultures grown in LB broth into LB 1% agar plates. Tips were stabbed perpendicular to the agar to the bottom of the plate. Plates were incubated at 37°C for 48 hours then the agar was removed, and plates were dried 10 minutes then stained with 1% crystal violet for 10 minutes. Plates were then washed three times with water and motility was quantified by measuring the radius from the point of inoculation. Swimming motility was assessed by stabbing 10 uL pipette tips dipped into overnight cultures grown in LB broth into LB 0.3% agar plates. Tips were stabbed perpendicular to the agar halfway down the plate. Plates were incubated at 30°C for 24 hours then motility was quantified by measuring the radius of bacterial movement from the point of inoculation. Mucoidy was assessed by streaking isolates onto *Pseudomonas* Isolation Agar plates and incubating at 37°C for 24 hours then at room temperature for 48 hours, followed by visual assessment. Sheen was assessed by spotting 3 uL overnight cultures grown in LB onto LB 1.5% agar plates. Plates were incubated at 37°C for 24 hours then at room temperature for 24 hours, followed by visual assessment. Amino acid auxotrophy was assessed by diluting overnight cultures grown in LB 1:500 into M9 minimal media supplemented with 0.2% 1 M MgSO4 and 0.4% glucose with or without 0.5% casamino acids. Cultures were incubated at 37°C for 24 hours then bacteria were resuspended, and growth was assessed by measuring absorbance at 600nm. Mucoidy, sheen, and auxotrophy were categorized as binary phenotypes.

### 2.4 Antibiotic and phage susceptibility testing

Antibiotic susceptibility was determined by broth microdilution to determine the MIC of C/T, or using HardyDisks^TM^ (Hardy Diagnostics, Santa Maria, CA) following the CLSI M02 guidelines for testing *P. aeruginosa* utilizing antibiotic-saturated disks (26). Zones of inhibition (ZOI) were measured after 16-18 hours of incubation at 37°C. Susceptibility breakpoints were assigned from the CLSI M100 guidelines (27). Phage susceptibility testing was assessed with double-layer agar plaque assay. Briefly, bottom agar plates were prepared containing TSB with 1.5% agar, 1 mM CaCl_2_ and 1 mM MgCl_2_. 1 mL of overnight bacterial culture grown in TSB was pelleted and resuspended in 1 mL SM+ buffer (50 mM TrisCl pH 7.4, 100 mM NaCl, 8 mM MgSO_4_, 5 mM CaCl_2_) then diluted 1:10 in SM+ buffer. 130 uL of the dilution was mixed with 5 mL of molten top agar (TSB with 0.3% agar, 1 mM CaCl_2_ and 1 mM MgCl_2_) and then plated onto a bottom agar plate. After allowing the top agar to solidify, 5 uL of 10-fold serial dilutions of phage lysates were spotted on top of the plate and then allowed to dry. Plates were incubated upright at 37°C overnight, then the dilution with 10 or fewer individual plaques was used to determine the phage titer in the original lysate by calculating the plaque-forming units per mL (PFU/mL).

### 2.5 Statistical analysis

Differences in antibiotic and phage susceptibility were assessed with Wilcoxon Signed-Rank Test calculated using GraphPad Prism v10.6.1. PCA plots were created in RStudio.

### 2.6 Data availability

Assembled bacterial genomes were submitted to the National Center for Biotechnology Information (NCBI) under BioProject PRJNA1519935, with accession numbers listed in **Table S1**.

## 3. Results

### 3.1 Genomic and phenotypic diversity of *P. aeruginosa* collected before and after treatment-emergent resistance to ceftolozane-tazobactam

Paired *P. aeruginosa* clinical isolates from 28 unique patients (n=10 pwCF, n=18 non-pwCF) collected at baseline (pre-exposure) and following treatment-emergent resistance to ceftolozane-tazobactam (post-exposure, defined as ≥4-fold increase in C/T MIC compared to baseline) were included in this study. All patients were treated at UPMC for multidrug-resistant *P. aeruginosa* infections between 2016 −2024. A phylogenetic tree constructed from a core-genome alignment of all 56 isolates demonstrated that isolate pairs from each patient were more genetically similar to each other than to isolates from different patients (**Figure 1A**). All isolates were screened for *in vitro* phenotypes associated with acute and chronic virulence of *P. aeruginosa* (28–30). While differences in protease production, twitching motility, and amino acid autotrophy were observed when comparing CF to non-CF isolates, no differences were observed between isolates collected pre-versus post-exposure to C/T (**Figure S1**). This was further reflected in a principal component analysis of the virulence phenotype data, which showed some separation between the CF and non-CF isolate groups, while pre-and post-exposure isolate groups were almost entirely overlapping (**Figure 1B**). Together, these data suggest that while isolates from pwCF were phenotypically different from isolates from non-pwCF, pre-and post-exposure isolates were largely similar to one another both genomically and phenotypically.

**Figure 1.**
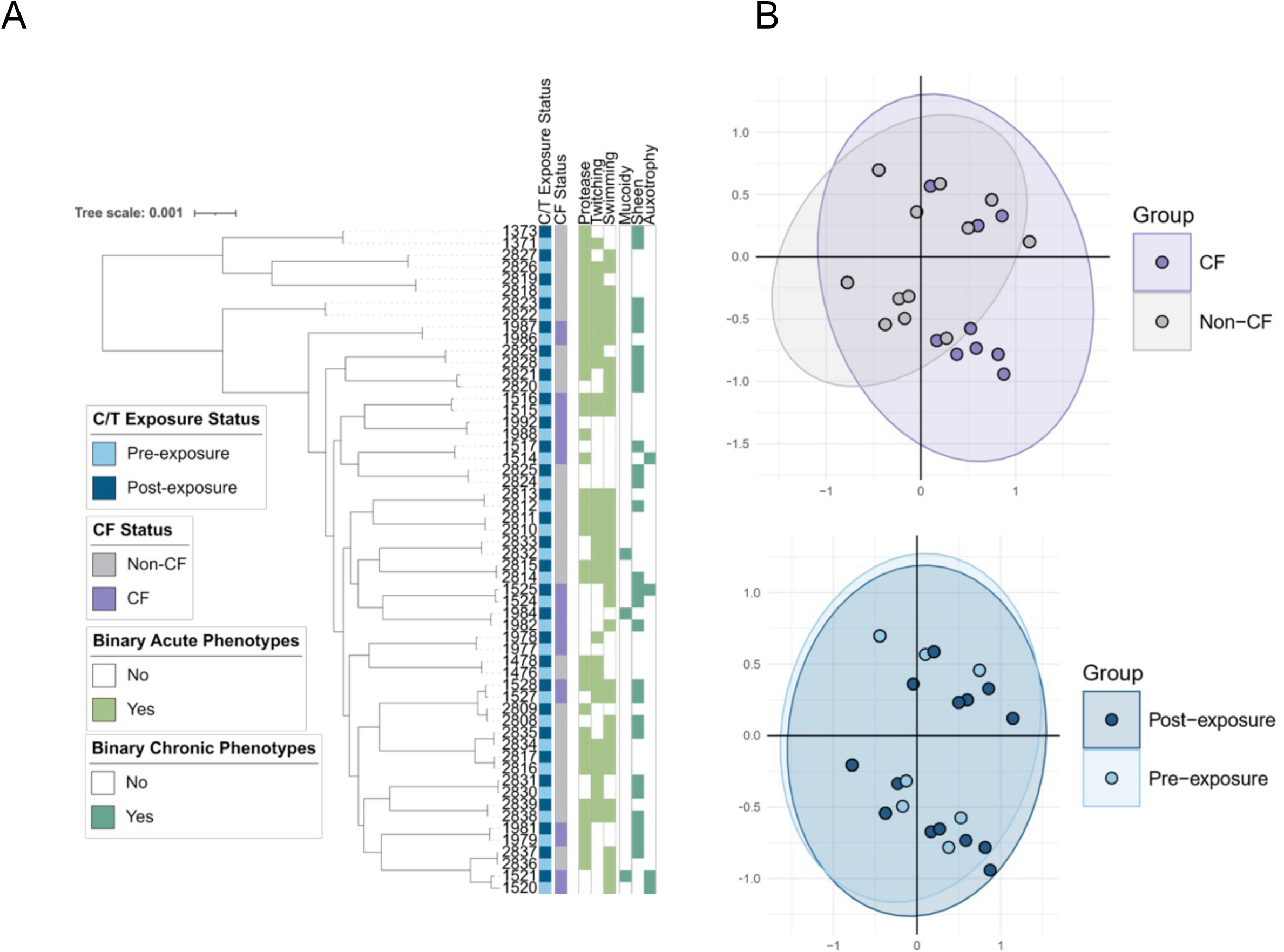
Genomic diversity and phenotypic profiling of *P. aeruginosa* clinical isolates collected at baseline and following treatment-emergent resistance to ceftolozane/tazobactam. (A) Phylogenetic tree showing genetic relatedness between pre-exposure and post-exposure isolate pairs across 28 unique patients (10 pwCF, 18 non-pwCF). Acute and chronic *in vitro* virulence phenotypes are shown for each isolate to the right of the tree. (B) Principal component analysis demonstrating the variance in binary coded expression of three acute phenotypes (protease, twitching motility, and swimming motility) and three chronic phenotypes (mucoidy, sheen, and amino acid auxotrophy) measured between isolates from pwCF and non-pwCF (top) and between pre-and post-C/T exposure isolates (bottom).

### 3.2 Treatment-emergent C/T resistance is associated with variable susceptibility to other antibiotics and phages

To determine whether treatment-emergent resistance to C/T was associated with altered susceptibility to other antibiotics, we performed Kirby Bauer disk diffusion assays on all 56 isolates against ten clinically relevant antibiotics belonging to multiple classes (**Figure 2**). Consistent with their inclusion in the study, all post-C/T exposure isolates had higher C/T MICs compared with the baseline isolate from the same patient. Among isolates collected from pwCF, there were no significant differences in susceptible to any other antibiotics between pre-and post-exposure isolates (**Figure 2A**). Among isolates collected from non-pwCF, post-exposure isolates were significantly less susceptible to both ceftazidime and ceftazidime/avibactam (**Figure 2B**), but there were no differences in susceptibility to any of the other antibiotics tested.

**Figure 2.**
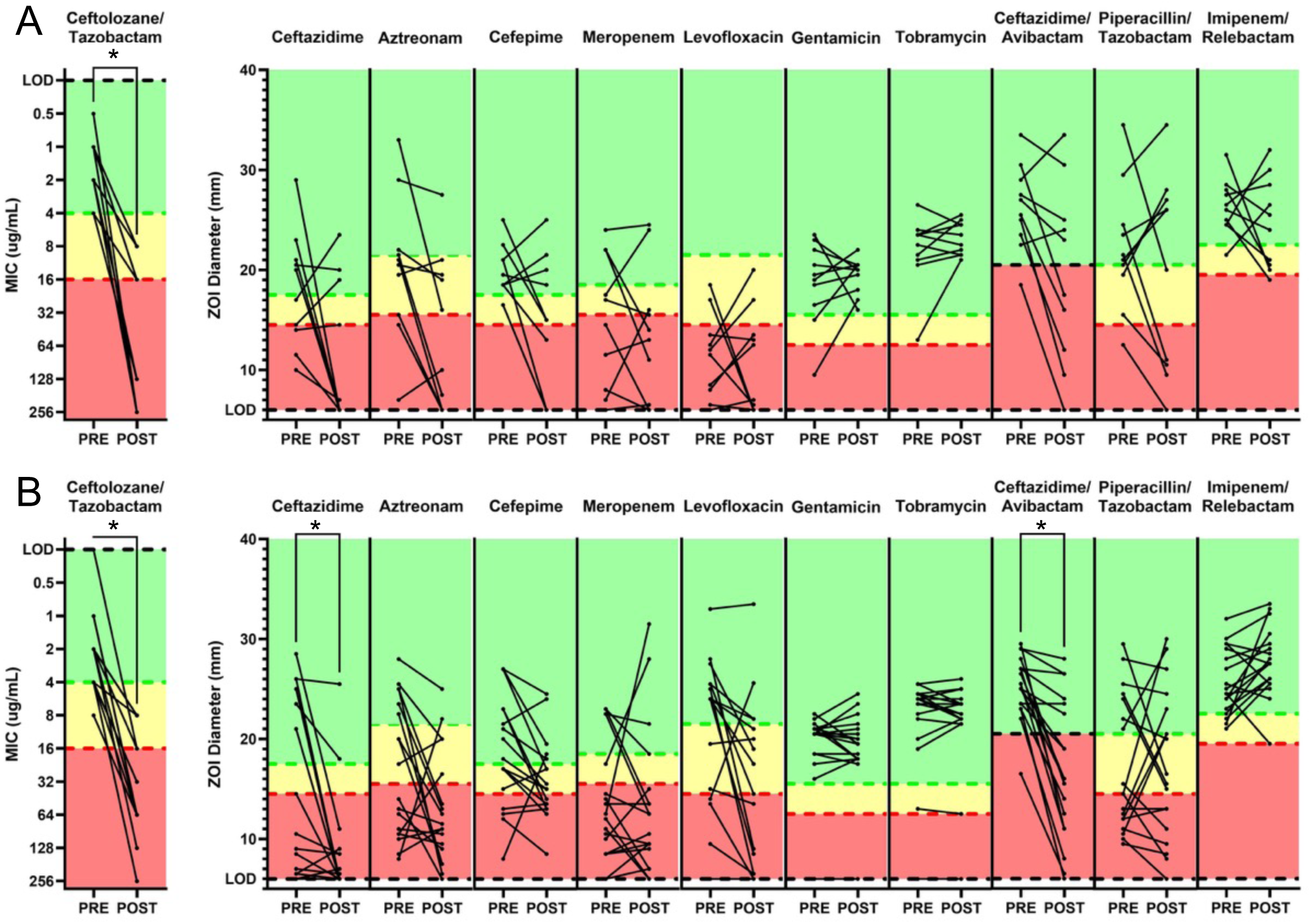
Differences in antibiotic susceptibility among pre-and post-C/T exposure *P. aeruginosa* isolates, separated by patient CF status. (A) Antibiotic susceptibility of pre-and post-exposure isolates from pwCF. (B) Antibiotic susceptibility of pre-and post-exposure isolates from non-pwCF. In both panels, C/T minimum inhibitory concentrations (MICs) are shown to the left; all other antibiotics were assessed via Kirby Bauer disk diffusion assay to determine the zone of inhibition (ZOI) diameter, in mm. Susceptibility cut-offs corresponding to susceptible (green), intermediate (yellow) and resistant (red) are shown for each antibiotic. Significant differences between pre-and post-exposure isolates were assessed with Wilcoxon Signed-Rank Test. * p < 0.05.

To determine whether treatment-emergent resistance to C/T was associated with altered susceptibility to *P. aeruginosa*-targeting bacteriophages, we tested the susceptibility of each isolate to seven genetically diverse lytic phages, some of which were described previously (2). When we compared phage titers between pre-and post-exposure isolates, most comparisons showed no change in phage susceptibility, though there were some modest trends toward increased susceptibility to phage PSA13 and decreased susceptibility to phages PSA07 and PSA62 among post-C/T exposure isolates (**Figure 3A**). We next assessed whether the number of active phages changed between same-patient pre-and post-exposure isolates collected from pwCF and non-pwCF and again observed no significant differences (**Figure 3B**). When we assessed changes in phage susceptibility by CF status and by phage, we found that isolates from pwCF were less susceptible to phages PSA56, PSA64, and PSA66 compared with isolates from non-pwCF (**Figure S2**). When we assess differences by patient, we observed 15 instances of post-exposure isolates gaining phage susceptibility and 15 instances of post-exposure isolates losing susceptibility (**Figure 3C**). While treatment-emergent C/T resistance did not correlate with greater phage susceptibility, 82% of the C/T-resistant isolates tested were susceptible to at least one phage included in the panel. Together these data suggest that while treatment-emergent C/T resistance did not result in predictable changes in phage susceptibility, phage therapy could still serve as a potential therapeutic option even in the presence of C/T resistance.

**Figure 3.**
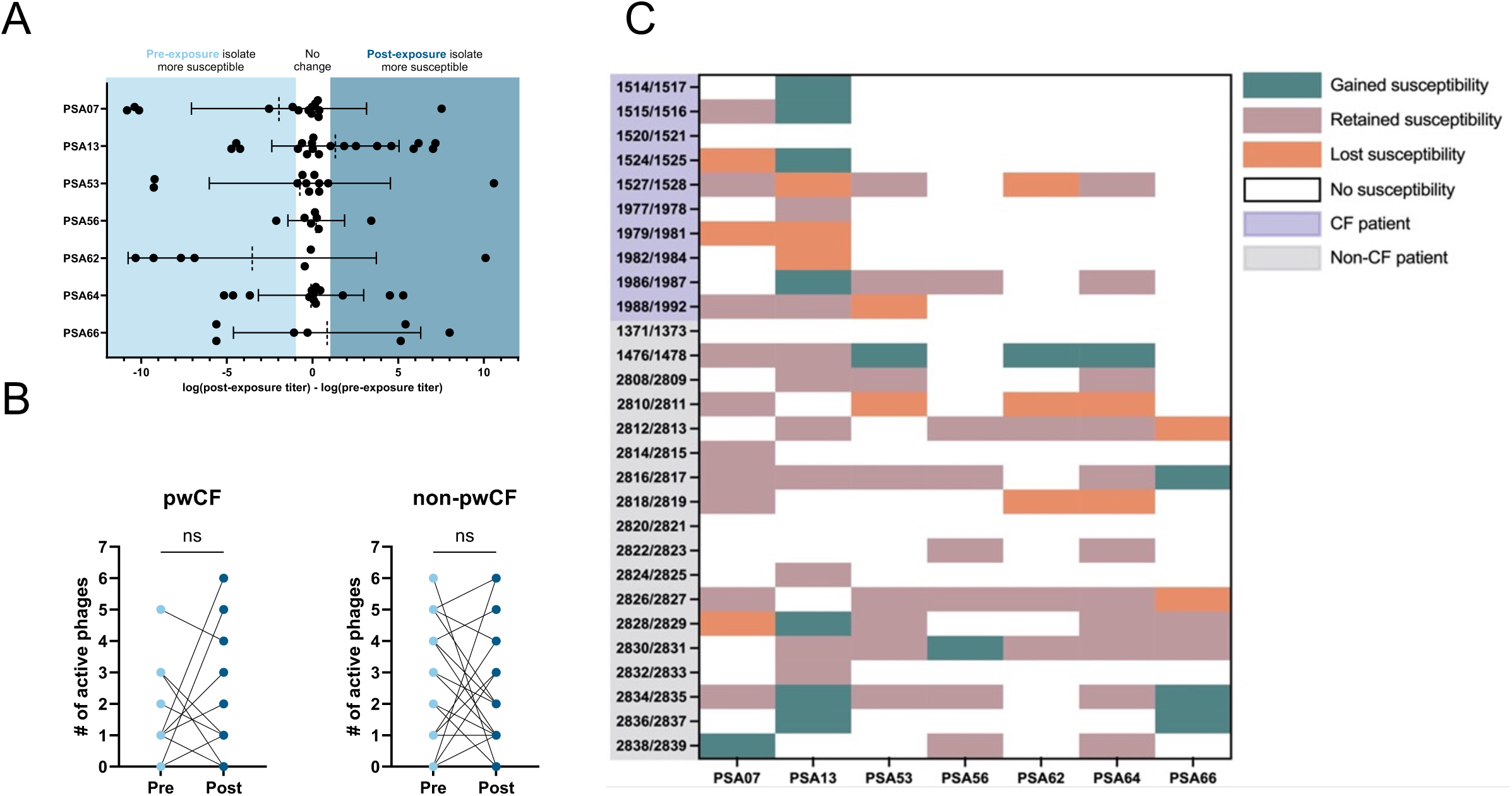
Differences in phage susceptibility between *P. aeruginosa* isolates. (A) Titer differences between isolates based on C/T exposure status. The log_10_ titer for each phage against each post-C/T exposure isolate was subtracted from the log_10_ titer against the pre-C/T exposure isolate from each patient. Dashed lines indicate mean and error bars show the standard deviation of all titer differences. Isolate pairs for which neither isolate was phage-susceptible are omitted. (B) Number of active phages against each isolate comparing pre-vs. post-C/T exposure status in pwCF (left) and non-pwCF (right). Lines connect values from the same patient. Significance was assessed with a Wilcoxon Signed Rank Test. (C) Changes in phage susceptibility between pre-and post-C/T exposure for isolate pairs from each patient. “Gained susceptibility” means the post-exposure isolate was more phage-susceptible; “Retained susceptibility” means the post-exposure isolate remained phage-susceptible; “Lost susceptibility” means the post-exposure isolate was less phage-susceptible; “No susceptibility” means both isolates were phage-resistant. Isolate pairs are shaded based on patient CF status.

### 3.3 Genetic Mutations Associated with Development of C/T Resistance

To determine underlying genetic factors contributing to treatment-emergent C/T resistance, the genotypes of five loci previously associated with C/T resistance were assessed to identify the emergence of variants in post-exposure isolates. These loci included AmpC (β-lactamase), AmpD (negative regulator of AmpC), FtsI (penicillin-binding protein 3), DacB (penicillin-binding protein 4), and MexRAB/OprM (efflux pump) (5,13,21). Among the five loci, AmpC mutations were most often associated with treatment-emergent C/T resistance, with 19/28 (68%) post-exposure isolates acquiring an AmpC mutation (**Figure 4A**). Treatment-emergent resistance-associated mutations were also observed in AmpD, FtsI, DacB, and MexR/MexB, with most isolates acquiring unique mutations at these loci. Five post-exposure isolates (18%) had no mutations detected in any of the five loci, and these isolates generally had lower fold-increases in C/T MIC compared with the other isolates. To assess the impact of mutations at each locus individually, we compared the C/T MICs of all isolates (both pre-and post-exposure) based on their genotype at each locus, and observed independent associations between mutations in AmpC, FtsI, DacB, and MexR/MexB and higher C/T MICs (**Figure 4B**). These data suggest that while C/T resistance can be driven by a variety of mutations at different loci, AmpC mutations appear to play a dominant role.

**Figure 4.**
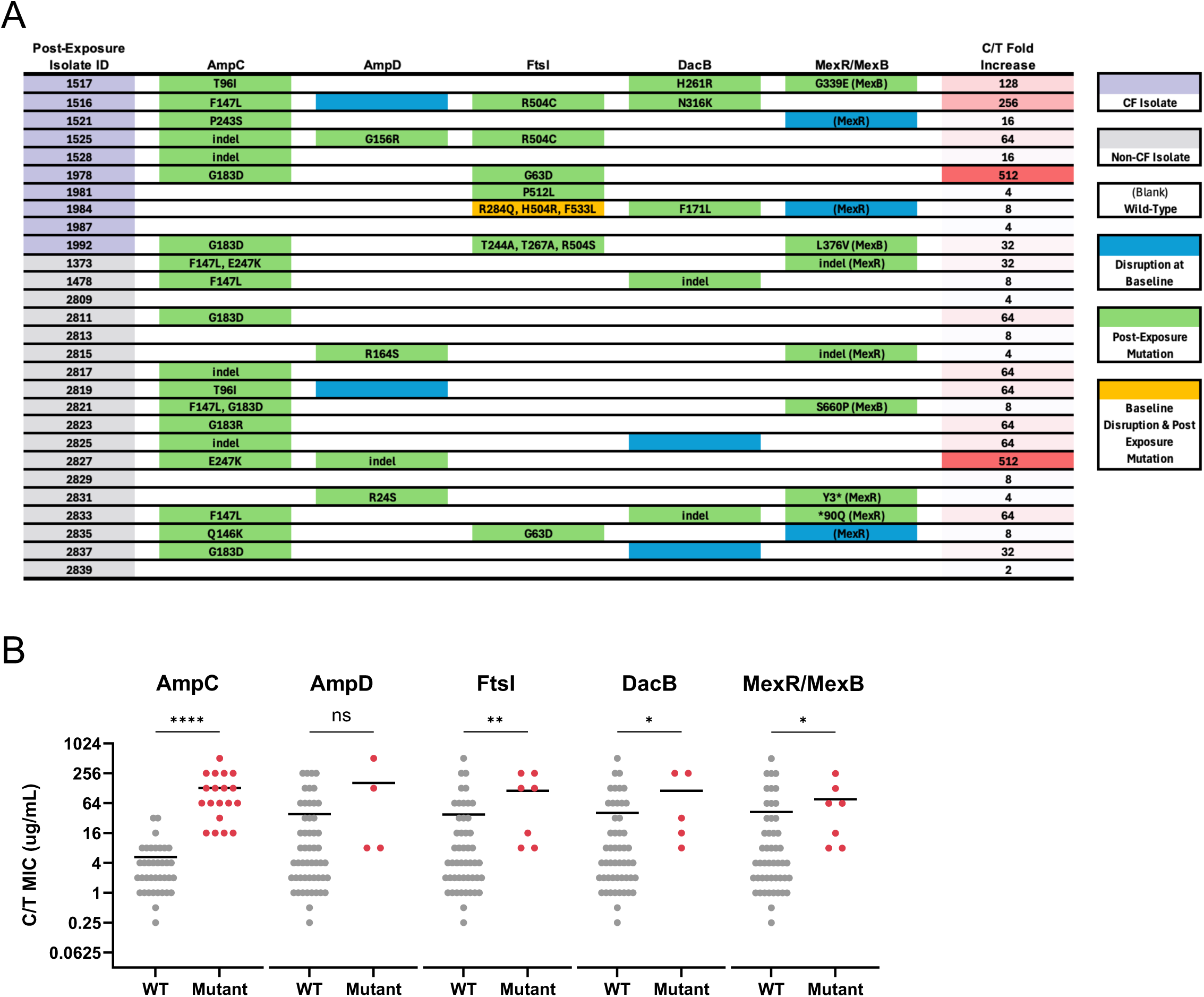
Mutations associated with treatment-emergent C/T resistance. (A) Mutations identified in post-exposure C/T-resistant isolates in AmpC, AmpD, FtsI, DacB, and MexR/MexB. Each mutation is annotated with the amino acid change. Indel = insertion/deletion variant. (B) C/T MICs comparing isolates that have wild-type (WT) or mutant genotypes for each locus.

## 4. Discussion

*P. aeruginosa* is an opportunistic pathogen that causes pneumonia and bacteremia in pwCF. The increasing prevalence of MDR *P. aeruginosa* limits our clinical arsenal of effective antibiotics for treating these infections. Ceftolozane-tazobactam (C/T) is a front-line antibiotic for the treatment of MDR *P. aeruginosa* infections. Despite promising clinical utility, resistance to C/T has been observed, further prompting the need to develop novel antimicrobials, such as phage therapy. To inform these development efforts, here we investigated the collateral effects of evolved C/T resistance on virulence-associated phenotypes, antibiotic susceptibility, and phage susceptibility among *P. aeruginosa* collected from both pwCF and non-pwCF. Our results suggest that virulence and antimicrobial phenotypes are variable between isolates, and do not change in predictable ways following treatment-emergent resistance to C/T.

Whole genome sequencing of *P. aeruginosa* clinical isolates collected from 28 patients before and after treatment-emergent resistance to C/T showed that isolates from the same patient, regardless of CF status, were more genetically related to one another than they were to isolates from different patients. This is consistent with prior reports, which have shown that bacterial isolates collected from the same patient are usually more closely related to each other compared to isolates from different patients (31,32). While both acute and chronic *in vitro* virulence phenotypes were also similar between pre-exposure and post-exposure isolates, we observed some differences in these phenotypes depending on whether isolates were collected from pwCF versus non-pwCF. In particular, protease production and twitching motility were less prevalent among isolates from pwCF compared with non-pwCF, while amino acid auxotrophy was observed more frequently among isolates from pwCF. These data are consistent with *P. aeruginosa* isolates from pwCF downregulating acute virulence phenotypes while upregulating chronic phenotypes, which facilitate adaptation to the host environment in the face of host-associated selective pressures (33–35). We observed no differences in these phenotypes associated with treatment-emergent C/T resistance, suggesting that resistance emergence does not impact acute or chronic virulence in predictable ways.

To determine if C/T resistance mediates collateral resistance or susceptibility to other antimicrobials such as antibiotics and phages, we conducted susceptibility testing against C/T plus ten additional antibiotics and seven lytic phages for all 56 isolates in this study. When we compared the antibiotic susceptibilities of paired pre-and post-exposure isolates, we observed that while all post-exposure isolates were more resistant to C/T, consistent changes in resistance or susceptibility patterns for the other antibiotics were minimal. The only significant difference we observed was that among isolates from non-pwCF, post-exposure isolates were more resistant to ceftazidime and ceftazidime/avibactam. This finding is consistent with other studies, which have found cross-resistance to ceftazidime-avibactam among C/T-resistant isolates. Previous studies have also found associations between C/T resistance and increased sensitivity to piperacillin-tazobactam and imipenem-relebactam. While we did not observe a statistically significant shift in sensitivity towards these antibiotics, a large proportion of isolates, both pre-and post-C/T treatment, were susceptible to both agents (5,36–38). When we examined changes in phage susceptibility, no obvious trends were observed when comparing pre-versus post-C/T exposure isolates, however the phages PSA56, PSA64, and PSA66 exhibited significantly more activity on isolates from non-pwCF compared to isolates from pwCF. While the surface receptors of these phages are not fully characterized, we suspect they might use type-IV pili as receptors for adsorption to the bacterial cell. This would align with our finding that many isolates from pwCF do not display twitching motility, which is often associated with type-IV pili.

To explore genetic factors associated with treatment-evolved C/T resistance, we sequenced the genomes of all 56 clinical isolates and identified mutations in six candidate genes commonly associated with C/T resistance (5,21,39,40). Among the candidate genes examined, the greatest number of mutations were observed in the *ampC* gene, and isolates encoding *ampC* mutations had significantly higher C/T MICs compared to wild-type isolates. Furthermore, the highest fold increases in C/T MIC occurred in isolates with mutations in *ampC* plus one or more mutations in other candidate genes. These observations reinforce the idea that *ampC* mutations primarily contribute to treatment-emergent C/T resistance, but mutations in other genes can play a secondary role.

This study had several limitations. First, we studied a relatively small number of patients infected with *P. aeruginosa* and treated with a single antibiotic, thus the broader impacts of C/T and other antibiotic selection on phage susceptibility in both *P. aeruginosa* and other bacterial species warrant exploration. Second, we did not examine simultaneous application of phages and antibiotics, which might be a relevant exercise as phages often serve as adjunct therapy alongside small molecule antibiotics. Lastly, while we focused on the collateral impacts of treatment-emergent C/T resistance, it is unclear if the same trends (or lack thereof) that we observed would hold if we instead focused on treatment-emergent phage resistance. This will be a focus of our future work in this area.

In conclusion, we found that *P. aeruginosa* clinical isolates that display treatment-emergent resistance to C/T are diverse in their genotypes, phenotypes, and antibiotic and phage susceptibilities. Evolution in response to C/T exposure has the potential to increase resistance to other beta-lactam antibiotics, but the impact on phage susceptibility appears to be highly variable. While we did not find strong correlations between treatment-emergent C/T resistance and phage susceptibility, over 80% of post-exposure isolates were susceptible to at least one phage tested, suggesting that phage therapy could be useful for treating *P. aeruginosa* infections, including those that are C/T-resistant. This insight could guide the clinical application of next-generation antimicrobials for MDR pathogens.

---

**Figure S1.**
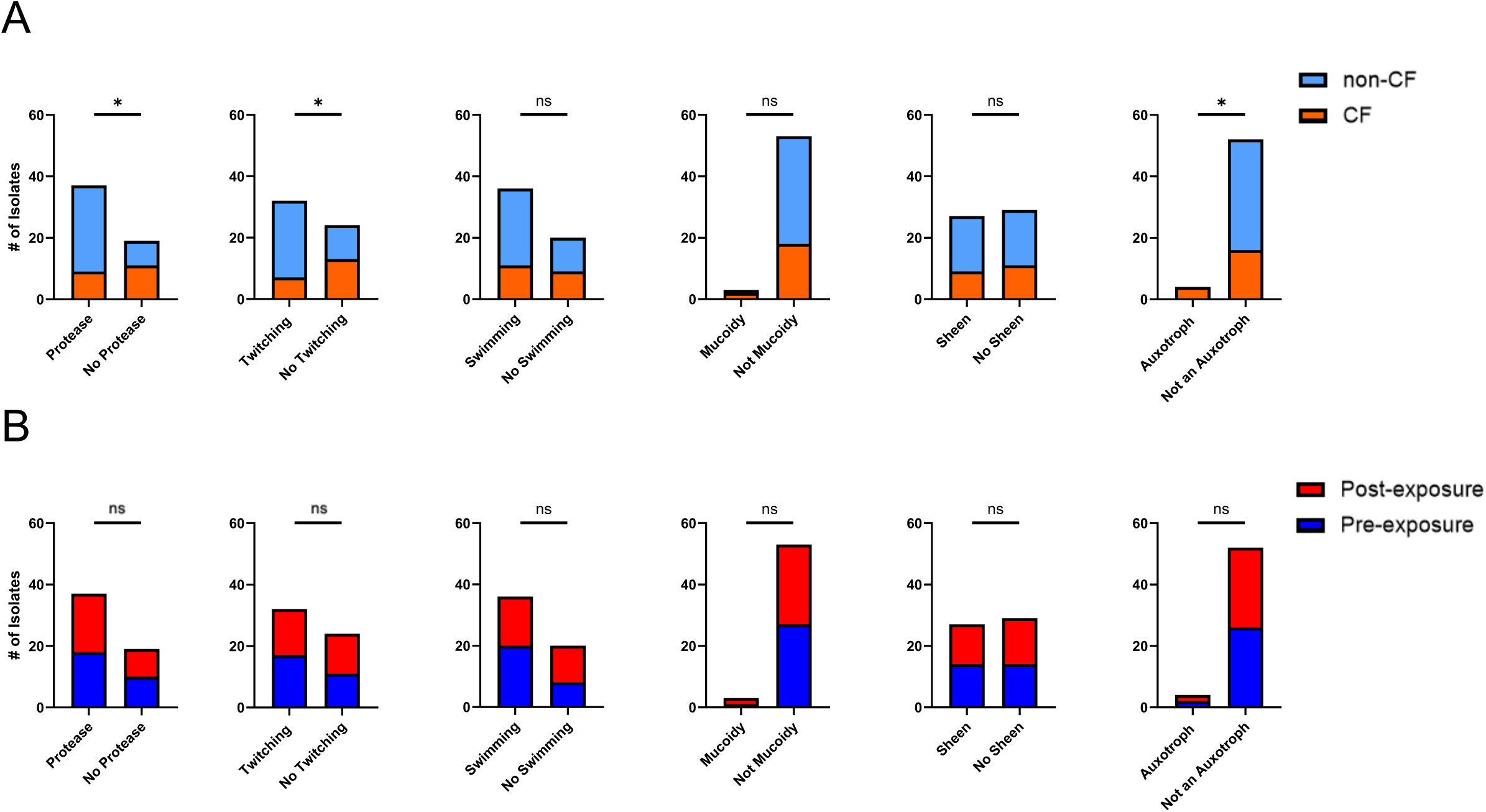
Acute and chronic phenotyping assays performed on all isolates. Protease, twitching and swimming were measured as a binary (0 = no, 1 = yes). (A) Comparison of non-CF versus CF isolates. (B) Comparison of C/T post-exposure and pre-exposure isolates.

**Figure S2.**
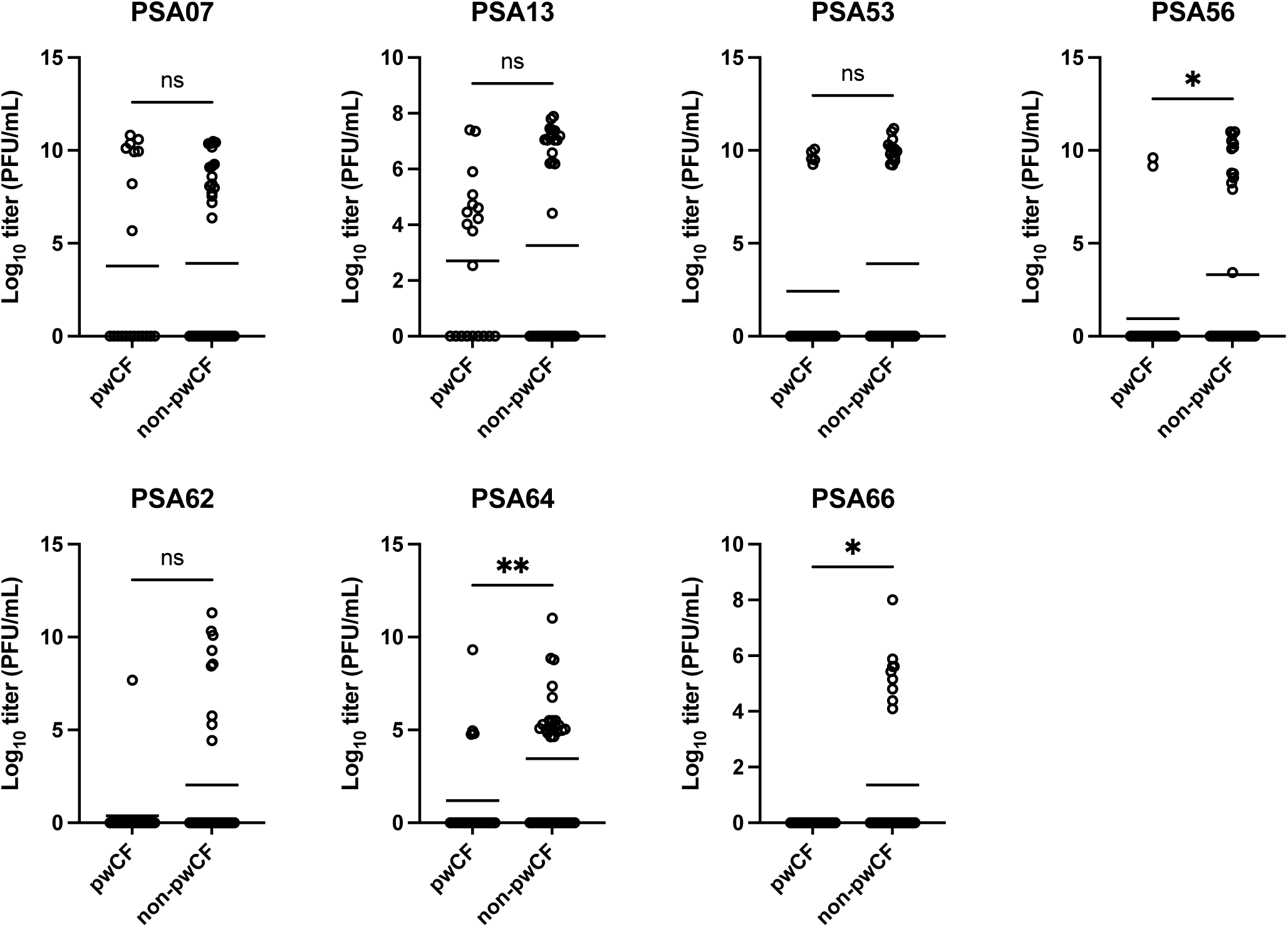
Phage susceptibility among all isolates collected from pwCF versus non-pwCF. Log_10_ titer of each phage against each isolate tested is shown. Phage-resistant isolates (i.e., isolates with no titer) are plotted at log_10_ titer = 0. Horizontal lines show the mean value for each group. Groups were compared with a Wilcoxon rank-sum test. *p< 0.05, **p< 0.005.

## References

1. Friedman ND, Temkin E, Carmeli Y. The negative impact of antibiotic resistance. Clin Microbiol Infect. 2016 May;22(5):416–22. doi:10.1016/j.cmi.2015.12.002

2. Nordstrom HR, Evans DR, Finney AG, Westbrook KJ, Zamora PF, Hofstaedter CE, et al. Genomic characterization of lytic bacteriophages targeting genetically diverse Pseudomonas aeruginosa clinical isolates. iScience. 2022 Jun;25(6):104372. doi:10.1016/j.isci.2022.104372

3. Domingo-Calap P, Delgado-Martinez J. Bacteriophages: Protagonists of a Post-Antibiotic Era. Antibiotics. 2018 Jul 27;7(3):66. doi:10.3390/antibiotics7030066

4. Centers for Disease Control and Prevention (U.S.). Antibiotic resistance threats in the United States, 2019 [Internet]. Centers for Disease Control and Prevention (U.S.); 2019 Nov. Available from: https://stacks.cdc.gov/view/cdc/82532 doi:10.15620/cdc:82532

5. Rubio AM, Kline EG, Jones CE, Chen L, Kreiswirth BN, Nguyen MH, et al. *In* Vitro Susceptibility of Multidrug-Resistant Pseudomonas aeruginosa following Treatment-Emergent Resistance to Ceftolozane-Tazobactam. Antimicrob Agents Chemother. 2021 May 18;65(6):e00084–21. doi:10.1128/AAC.00084-21

6. Yin C, Alam MZ, Fallon JT, Huang W. Advances in Development of Novel Therapeutic Strategies against Multi-Drug Resistant Pseudomonas aeruginosa. Antibiotics. 2024 Jan 25;13(2):119. doi:10.3390/antibiotics13020119

7. Bonyadi P, Saleh NT, Dehghani M, Yamini M, Amini K. Prevalence of antibiotic resistance of Pseudomonas aeruginosa in cystic fibrosis infection: A systematic review and meta-analysis. Microb Pathog. 2022 Apr;165:105461. doi:10.1016/j.micpath.2022.105461

8. Crull MR, Somayaji R, Ramos KJ, Caldwell E, Mayer-Hamblett N, Aitken ML, et al. Changing Rates of Chronic Pseudomonas aeruginosa Infections in Cystic Fibrosis: A Population-Based Cohort Study. Clin Infect Dis. 2018 Sep 14;67(7):1089–95. doi:10.1093/cid/ciy215

9. Jackson L, Waters V. Factors influencing the acquisition and eradication of early Pseudomonas aeruginosa infection in cystic fibrosis. J Cyst Fibros. 2021 Jan;20(1):8–16. doi:10.1016/j.jcf.2020.10.008

10. Guillaume O, Butnarasu C, Visentin S, Reimhult E. Interplay between biofilm microenvironment and pathogenicity of Pseudomonas aeruginosa in cystic fibrosis lung chronic infection. Biofilm. 2022 Dec;4:100089. doi:10.1016/j.bioflm.2022.100089

11. Pang Z, Raudonis R, Glick BR, Lin TJ, Cheng Z. Antibiotic resistance in Pseudomonas aeruginosa: mechanisms and alternative therapeutic strategies. Biotechnol Adv. 2019 Jan;37(1):177–92. doi:10.1016/j.biotechadv.2018.11.013

12. Montaner M, Lopez-Arguello S, Oliver A, Moya B. PBP Target Profiling by β-Lactam and β-Lactamase Inhibitors in Intact Pseudomonas aeruginosa: Effects of the Intrinsic and Acquired Resistance Determinants on the Periplasmic Drug Availability. Papp-Wallace KM, editor. Microbiol Spectr. 2023 Feb 14;11(1):e03038–22. doi:10.1128/spectrum.03038-22

13. Cabot G, Bruchmann S, Mulet X, Zamorano L, Moya B e, Juan C, et al. Pseudomonas aeruginosa Ceftolozane-Tazobactam Resistance Development Requires Multiple Mutations Leading to Overexpression and Structural Modification of AmpC. Antimicrob Agents Chemother. 2014 May 14;58(6):3091–9. doi:10.1128/aac.02462-13

14. Fraile-Ribot PA, Cabot G, Mulet X, Perianez L, Martin-Pena ML, Juan C, et al. Mechanisms leading to in vivo ceftolozane/tazobactam resistance development during the treatment of infections caused by MDR Pseudomonas aeruginosa. J Antimicrob Chemother. 2018 Mar 1;73(3):658–63. doi:10.1093/jac/dkx424

15. Haidar G, Philips NJ, Shields RK, Snyder D, Cheng S, Potoski BA, et al. Ceftolozane-Tazobactam for the Treatment of Multidrug-Resistant Pseudomonas aeruginosa Infections: Clinical Effectiveness and Evolution of Resistance. Clin Infect Dis Off Publ Infect Dis Soc Am. 2017 Jul 1;65(1):110–20. doi:10.1093/cid/cix182

16. Gomis-Font Mia A, Cabot G, Sanchez-Diener I, Fraile-Ribot PA, Juan C, Moya B, et al. In vitro dynamics and mechanisms of resistance development to imipenem and imipenem/relebactam in Pseudomonas aeruginosa. J Antimicrob Chemother. 2020 Sep 1;75(9):2508–15. doi:10.1093/jac/dkaa206

17. Lyon R, Jones RA, Shropshire H, Aberdeen I, Scanlan DJ, Millard A, et al. Membrane lipid renovation in *Pseudomonas* aeruginosa implications for phage therapy? Environ Microbiol. 2022 Oct;24(10):4533–46. doi:10.1111/1462-2920.16136

18. El-Shibiny A, El-Sahhar S. Bacteriophages: the possible solution to treat infections caused by pathogenic bacteria. Can J Microbiol. 2017 Nov;63(11):865–79. doi:10.1139/cjm-2017-0030

19. Chan BK, Stanley GL, Kortright KE, Vill AC, Modak M, Ott IM, et al. Personalized inhaled bacteriophage therapy for treatment of multidrug-resistant Pseudomonas aeruginosa in cystic fibrosis. Nat Med. 2025 May;31(5):1494–501. doi:10.1038/s41591-025-03678-8

20. Terlizzi V, Rinninella G, Viglietto L, D’Agosto A, Di Luca M, Lopes-Pacheco Meias, et al. Phage therapy in people with cystic fibrosis: A systematic review. Int J Infect Dis. 2026 Jun;167:108581. doi:10.1016/j.ijid.2026.108581

21. Shah S, Kline EG, Haidar G, Squires KM, Pogue JM, McCreary EK, et al. Rates of Resistance to Ceftazidime-Avibactam and Ceftolozane-Tazobactam Among Patients Treated for Multidrug-Resistant Pseudomonas aeruginosa Bacteremia or Pneumonia. Clin Infect Dis. 2025 Jan 15;80(1):24–8. doi:10.1093/cid/ciae332

22. Wick RR, Judd LM, Gorrie CL, Holt KE. Unicycler: Resolving bacterial genome assemblies from short and long sequencing reads. PLoS Comput Biol. 2017 Jun;13(6):e1005595. doi:10.1371/journal.pcbi.1005595

23. Page AJ, Cummins CA, Hunt M, Wong VK, Reuter S, Holden MTG, et al. Roary: rapid large-scale prokaryote pan genome analysis. Bioinformatics. 2015 Nov 15;31(22):3691–3. doi:10.1093/bioinformatics/btv421

24. Seemann T. Prokka: rapid prokaryotic genome annotation. Bioinformatics. 2014 Jul 15;30(14):2068–9. doi:10.1093/bioinformatics/btu153

25. Stamatakis A. RAxML version 8: a tool for phylogenetic analysis and post-analysis of large phylogenies. Bioinformatics. 2014 May 1;30(9):1312–3. doi:10.1093/bioinformatics/btu033

26. CLSI. Performance Standards for Antimicrobial Disk Susceptibility Tests. 13th ed. CLSI standard M02. 2018.

27. CLSI. Performance Standards for Antimicrobial Susceptibility Testing. 33rd ed. CLSI supplement M100. 2023.

28. Flores-Vega Vonica R, Hernandez-Martinez G, Cocotl-Yanez M, Ares MA, Lincopan N, Ortiz-Navarrete V, et al. Contribution of two-component regulatory systems to the acute-to-chronic infection transition of Pseudomonas aeruginosa in cystic fibrosis. J Bacteriol. 2026 Feb 25;208(3):e00471–25. doi:10.1128/jb.00471-25

29. Hogardt M, Heesemann Jurgen. Adaptation of Pseudomonas aeruginosa during persistence in the cystic fibrosis lung. Int J Med Microbiol IJMM. 2010 Dec;300(8):557–62. doi:10.1016/j.ijmm.2010.08.008

30. Liao C, Huang X, Wang Q, Yao D, Lu W. Virulence Factors of Pseudomonas Aeruginosa and Antivirulence Strategies to Combat Its Drug Resistance. Front Cell Infect Microbiol. 2022 Jul 6;12:926758. doi:10.3389/fcimb.2022.926758

31. Persyn E, Sassi M, Aubry M, Broly M, Delanou S, Asehnoune K, et al. Rapid genetic and phenotypic changes in Pseudomonas aeruginosa clinical strains during ventilator-associated pneumonia. Sci Rep. 2019 Mar 18;9:4720. doi:10.1038/s41598-019-41201-5

32. Datar R, Coello Pelegrin A, Orenga S, Chalansonnet Verie, Mirande C, Dombrecht J, et al. Phenotypic and Genomic Variability of Serial Peri-Lung Transplantation Pseudomonas aeruginosa Isolates From Cystic Fibrosis Patients. Front Microbiol. 2021 Apr 7;12:604555. doi:10.3389/fmicb.2021.604555

33. Perez LRR, Costa MCN, Freitas ALP, Barth AL. Evaluation of biofilm production by Pseudomonas Aeruginosa isolates recovered from cystic fibrosis and non-cystic fibrosis patients. Braz J Microbiol. 2011;42(2):476–9. doi:10.1590/S1517-838220110002000011

34. Cullen L, Weiser R, Olszak T, Maldonado RF, Moreira AS, Slachmuylders L, et al. Phenotypic characterization of an international Pseudomonas aeruginosa reference panel: strains of cystic fibrosis (CF) origin show less in vivo virulence than non-CF strains. Microbiology. 2015;161(10):1961–77. doi:10.1099/mic.0.000155

35. Perez LRR, de Freitas ALucia P, Barth AL is. Cystic and Non-Cystic Fibrosis Pseudomonas aeruginosa Isolates are not Differentiated by the Quorum-Sensing Signaling and Biofilm Production. Curr Microbiol. 2012 Jan 1;64(1):81–4. doi:10.1007/s00284-011-0041-z

36. Fernandez-Esgueva M, Lopez-Calleja AI, Mulet X, Fraile-Ribot PA, Cabot G, Huarte R, et al. Characterization of AmpC β-lactamase mutations of extensively drug-resistant Pseudomonas aeruginosa isolates that develop resistance to ceftolozane/tazobactam during therapy. Enferm Infecc Microbiol Clin. 2020 Dec;38(10):474–8. doi:10.1016/j.eimc.2020.01.017

37. Skoglund E, Abodakpi H, Rios R, Diaz L, De La Cadena E, Dinh AQ, et al. In Vivo Resistance to Ceftolozane/Tazobactam in Pseudomonas aeruginosa Arising by AmpC-and Non-AmpC-Mediated Pathways. Case Rep Infect Dis. 2018 Dec 23;2018:9095203. doi:10.1155/2018/9095203

38. Rieper F, Wittmann J, Bunk B, Sproer C, Hafner M, Willy C, et al. Systematic bacteriophage selection for the lysis of multiple Pseudomonas aeruginosa strains. Front Cell Infect Microbiol. 2025 May 23;15. doi:10.3389/fcimb.2025.1597009

39. Mechanisms of Resistance to Ceftolozane/Tazobactam in Pseudomonas aeruginosa: Results of the GERPA Multicenter Study. Antimicrob Agents Chemother [Internet]. Available from: https://journals.asm.org/doi/10.1128/aac.01117-20

40. Nguyen HA, Peleg AY, Song J, Wisniewski JA, Blakeway LV, Badoordeen GZ, et al. Complex pathways to ceftolozane-tazobactam resistance in clinical *Pseudomonas* aeruginosa isolates: a genomic epidemiology study. Clin Microbiol Infect. 2026 Jan 1;32(1):110–7. doi:10.1016/j.cmi.2025.09.015

